# Seq2Topt: a sequence-based deep learning predictor of enzyme optimal temperature

**DOI:** 10.1101/2024.08.12.607600

**Authors:** Sizhe Qiu, Bozhen Hu, Jing Zhao, Weiren Xu, Aidong Yang

## Abstract

An accurate deep learning predictor is needed for enzyme optimal temperature (*T*_*opt*_), which quantitatively describes how temperature affects the enzyme catalytic activity. Seq2Topt, developed in this study, reached a superior accuracy on *T*_*opt*_ prediction just using protein sequences (RMSE = 13.3℃ and R2=0.48) in comparison with existing models, and could capture key protein regions for enzyme *T*_*opt*_ with multi-head attention on residues. Through case studies on thermophilic enzyme selection and predicting enzyme *T*_*opt*_ shifts caused by point mutations, Seq2Topt was demonstrated as a promising computational tool for enzyme mining and *in-silico* enzyme design. Additionally, accurate deep learning predictors of enzyme optimal pH (Seq2pHopt, RMSE=0.92 and R2=0.37) and melting temperature (Seq2Tm, RMSE=7.57℃ and R2=0.64) were developed based on the model architecture of Seq2Topt, suggesting that the development of Seq2Topt could potentially give rise to a useful prediction platform of enzymes.

## 1. Introduction

Temperature is an important influencing factor of enzyme catalysis [1], awnd researchers, especially of enzyme mining or engineering, want to quantitatively characterize the thermophilicity of enzymes, that is the enzyme optimal temperature (*T*_*opt*_). Given the large gap of enzyme *T*_*opt*_ in databases (e.g., BRENDA [2]) [3] and the high cost of enzyme assays [4], using machine learning models to predict enzyme *T*_*opt*_ has the potential to yield an optimal solution.

Most existing machine learning (ML) models of enzyme *T*_*opt*_ are specific to certain enzyme classes, such as Zhang and Ge, 2011’s model for xylanases [5]. Those models were usually developed on a small dataset specific to one class of enzymes, and the feature generation was mostly statistical descriptors of protein sequences (e.g., amino acid composition or dipeptide composition) [5–7]. Though some enzyme specific predictors had good accuracy (e.g., Chu et al., 2016’s model for beta-agarases [7] or Yan and Wu, 2019’s model for beta-glucosidases [8]), their restricted scope limited their applications. Therefore, a general predictor of *T*_*opt*_, regardless of enzyme classes, is needed by researchers who want to perform enzyme mining from massive sequencing data or computer aided engineering of enzymes.

Regarding *T*_*opt*_ prediction for general enzymes, there only exist two tools, TOMER [3][9] and Preoptem [10]. TOMER, an ensemble model, can accurately predict *T*_*opt*_ with a R2 score of 0.94, but it requires the optimal growth temperature (OGT) of the organism as an extra input other than the protein sequence. Also, the feature importance analysis of TOMER showed that the OGT contributed almost 50% of its predictive power [3], which might cause bias for the same enzymes expressed in different microorganisms. The strong reliance of TOMER on the OGT makes it inconvenient to use, because the OGT values are not accessible in many scenarios without the organismal information (e.g., enzyme mining from metagenomics or cell-free enzyme catalysis). In contrast, Preoptem, a deep learning model using one-hot encoding and CNN, can predict *T*_*opt*_ just from protein sequences. However, its prediction accuracy is not high (R2=0.36) and it cannot provide feature importance interpretation. In conclusion, there lacks an accurate predictive model of enzyme *T*_*opt*_ just using protein sequences.

With the aim to enhance the prediction accuracy of enzyme *T*_*opt*_ just from protein sequences, this study used a pre-trained language model of proteins, multi-head attention mechanism, and residual dense neural networks to construct a deep learning model, Seq2Topt, with good accuracy. Also, Seq2Topt allows the interpretation of attention weights on protein residues, which helps to decipher the key sequence information for enzyme *T*_*opt*_. Through case studies on the selection of thermophilic enzymes and predicting *T*_*opt*_ shifts caused by point mutations, this work demonstrated that Seq2Topt could function as a powerful computational tool to aid enzyme mining and engineering.

## 2. Methods

### 2.1 Datasets

For *T*_*opt*_, the dataset was obtained from the github repository of TOMER [9], which had 2917 entries. 10% of the dataset was randomly split as the holdout test set (n=291), and the remaining 90% of the dataset was used as the training set (n=2626). The holdout test set (https://github.com/SizheQiu/Seq2Topt/tree/main/data/Topt/test.csv) was used in the model comparison of TOMER, Seq2Topt and Preoptem. To mediate the imbalance of *T*_*opt*_ in the training dataset, oversampling was performed to double entries at high *T*_*opt*_ (≥80℃) by randomly duplicating existing entries at those value ranges (**see SI, Figure S1**). The training and test datasets of melting temperature (*T*_*m*_) (T≥50℃) were obtained from the github repository of DeepTM [11]. The training and test datasets of *T*_*m*_ had 6240 and 1550 entries, respectively. The training, validation and test datasets of optimal pH (*pH*_*opt*_) were obtained from the Zenodo repository of EpHod [12]. The training, validation and test datasets of *pH*_*opt*_ had 7124, 760, and 1971 entries, respectively.

### 2.2 Construction of the deep learning model

Seq2Topt consisted of sequence embedding computation by ESM-2 [13], multi-head attention (n_head=4) and 3 residual dense blocks (**Figure 1**). First, the protein sequence embeddings (*r* ∈ *R*^*L*∗*dim*^, *L*: *sequence lengt*ℎ, *dim* = 320) were computed using the esm2_t6_8M_UR50D model (https://github.com/facebookresearch/esm). The embeddings (*r*) were passed to a convolutional neural network (CNN) to generate the values (*v* ∈ *R*^*L*∗*dim*^), and a CNN and softmax to generate the weights (*w* ∈ *R*^*L*∗*dim*^). Then, element-wise products of values and weights were computed as the attention weighted features (*x*_*att*_ ∈ *R*^*dim*∗*dim*^). In the multi-head attention process, the attention weighted features were computed for 4 times, and all MaxPools (*x*^*k*^_*max*_ ∈ *R*^1∗*dim*^, *k*: *index of attention* ℎ*ead*) and Sums (*x*^*k*^_*sum*_ ∈ *R*^1∗*dim*^, *k*: *index of attention* ℎ*ead*) of attention weighted features were concatenated as the inputs (*x*_*concat*_ ∈ *R*^2∗4∗1280^) for residual dense blocks. Each residual dense block consisted of a linear layer, batch normalization, activation layer, and dropout layer. The activation function used in each residual dense block was Leaky ReLU [14]. The concatenated feature was passed through 3 residual dense blocks and a linear layer to regress for the target value.

**Figure 1.**
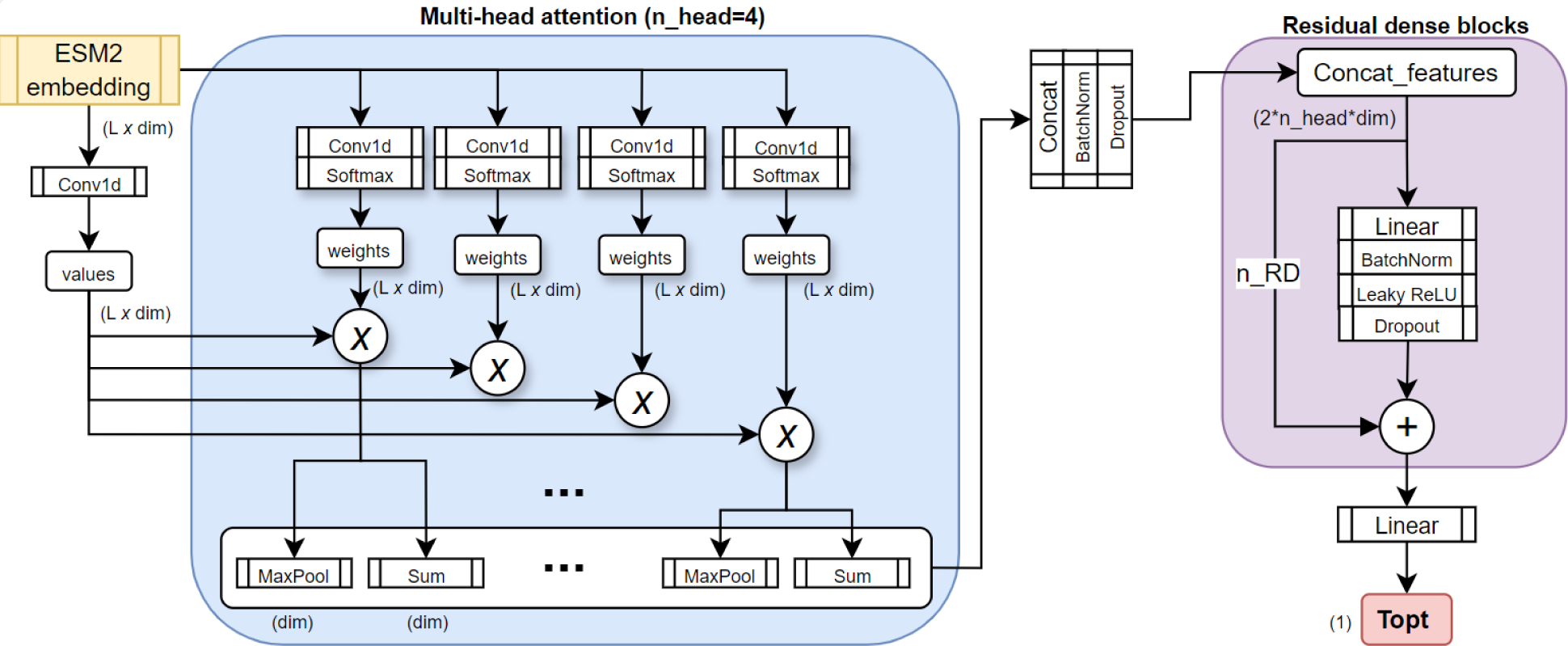
The model architecture of Seq2Topt. L: protein sequence length; dim: embedding dimension size; Conv1d: 1-D convolutional layer; Ⓧ: element-wise multiplication; BatchNorm: Batch normalization (re-centering and re-scaling); n_head: the number of heads in multi-head attention; RD: residual dense block, a dense layer with residual connection; n_RD=3; *T*_*opt*_: optimal temperature.

### 2.3 Deep learning model training

For the training process, batch training was used (batch size=32) for the efficiency and generalizability of the deep learning neural network. Adam optimization algorithm [15] was used to update neural network weights iteratively. The loss function was mean squared error (MSE). The initial learning rate was 0.0005, and the learning rate decayed by 50% for every 10 epochs to prevent overfitting. Before model training started, 10% of the training set was splitted out as the validation set, and target values were rescaled as 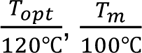, and 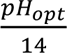. During the training process, the prediction accuracy of the model was evaluated with root mean squared error (RMSE), mean average error (MAE), and r-squared (R2) (**see SI, Supplementary methods section S1.2**). For details of software and hardware, please see section S1.1 of the supplementary information. The default hyperparameter was {window size:5, dropout rate: 0.1, number of attention heads: 4, number of residual dense blocks: 3}. The hyperparameter optimization was performed on the sliding window size of the CNN (3, 5, 7) and the dropout rate (0.1, 0.2, 0.5) (**see SI, Figure S2**).

### 2.4 Interpretation of residue attention weights

To investigate how enzyme *T*_*opt*_ was predicted from the amino acid sequence, the average attention weights on residues (*w*_*avg*_ ∈ *R*^*L*∗1^) were computed by averaging the weights (*w* ∈ *R*^*L*∗*dim*^) across the embedding feature dimension (dim), and across 4 attention heads. Then, the average residue attention weights (*w*_*avg*_) were mapped to the protein sequence, together with protein structural features (e.g., helix or turn) obtained from the uniprot database [16]. The spatial distribution of high attention weights and regions of annotated protein structural features could assist in revealing the hidden key sequence information influencing the enzyme ^*T*^_*opt*_.

## 3. Results

### 3.1 The superior performance of Seq2Topt

With the optimal set of hyperparameters (**see SI, Figure S2**), the training process reduced the RMSE and MAE to 13.3℃ and 10.5℃, and enhanced R2 from around −2.5 to 0.48 (**Figure 2AB, Figure S3**). For model comparison, this study used TOMER [3][9] and Preoptem [10] to predict *T*_*opt*_ values for the same holdout test set (**Figure 2CF**). Seq2Topt had a higher accuracy than Preoptem with lower RMSE and higher R2 scores, and a close accuracy to TOMER, which used both protein sequences and OGTs as inputs (**Figure 2DG**). Regarding the high temperature value range in the test set (>60℃), the prediction error of Seq2Topt was relatively smaller than that of Preoptem, but higher than that of TOMER. Overall, the accuracy assessment on *T*_*opt*_ prediction exhibited the superior performance of Seq2Topt.

**Figure 2.**
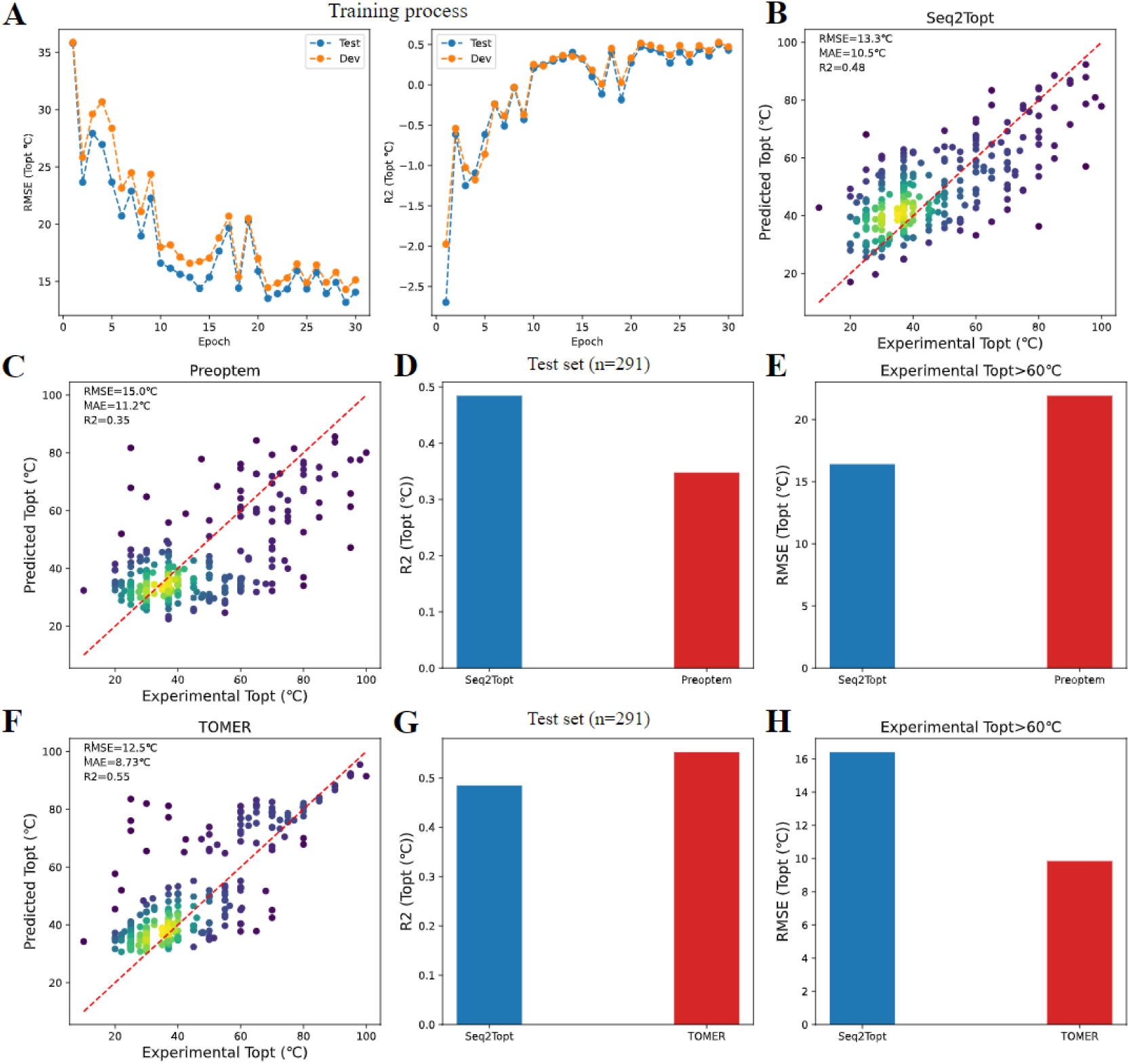
The assessment of the model performance. (A) The RMSE and R2 scores of *T*_*opt*_ prediction during the training process. (B) Experimental and predicted *T*_*opt*_ by Seq2Topt (RMSE=13.3℃ and R2=0.48). (C) Experimental and predicted *T*_*opt*_ by Preoptem (RMSE=15.0℃ and R2=0.35). (D) The comparison of R2 scores of *T*_*opt*_ prediction for the test set (R2 of Seq2Topt=0.48 and R2 of Preoptem=0.35). (E) The comparison of RMSE scores of *T*_*opt*_ prediction at the high temperature value range (RMSE of Seq2Topt=16.4℃ and RMSE of Preoptem=21.9℃). (F) Experimental and predicted *T*_*opt*_ by TOMER (RMSE=12.5℃ and R2=0.55). (G) The comparison of R2 scores of *T*_*opt*_ prediction for the test set (R2 of Seq2Topt=0.48 and R2 of TOMER=0.55). (H) The comparison of RMSE scores of *T*_*opt*_ prediction at the high temperature value range (RMSE of Seq2Topt=16.4℃ and RMSE of TOMER=9.8℃).

### 3.2 Using Seq2Topt to estimate the thermophilicity of organisms and enzymes

328 enzymes from representative mesophilic, thermophilic, and hyperthermophilic microorganisms (**see SI, Table S1**) were selected to examine the prediction performance of Seq2Topt on microorganisms and enzymes of different thermophilicities. First, the comparison between experimental and predicted *T*_*opt*_ values showed that Seq2Topt had good accuracy on enzymes from both mesophiles (RMSE=10.86℃, MAE=8.56℃) and thermophiles/hyperthermophiles ((RMSE=14.07℃, MAE=10.81℃) (**Figure 3AB**). The significant differential distributions of predicted *T*_*opt*_ values of enzymes from mesophiles and thermophiles/hyperthermophiles demonstrated that Seq2Topt could classify enzymes and microorganisms of different thermophilicities (**Figure 3C**).

**Figure 3.**
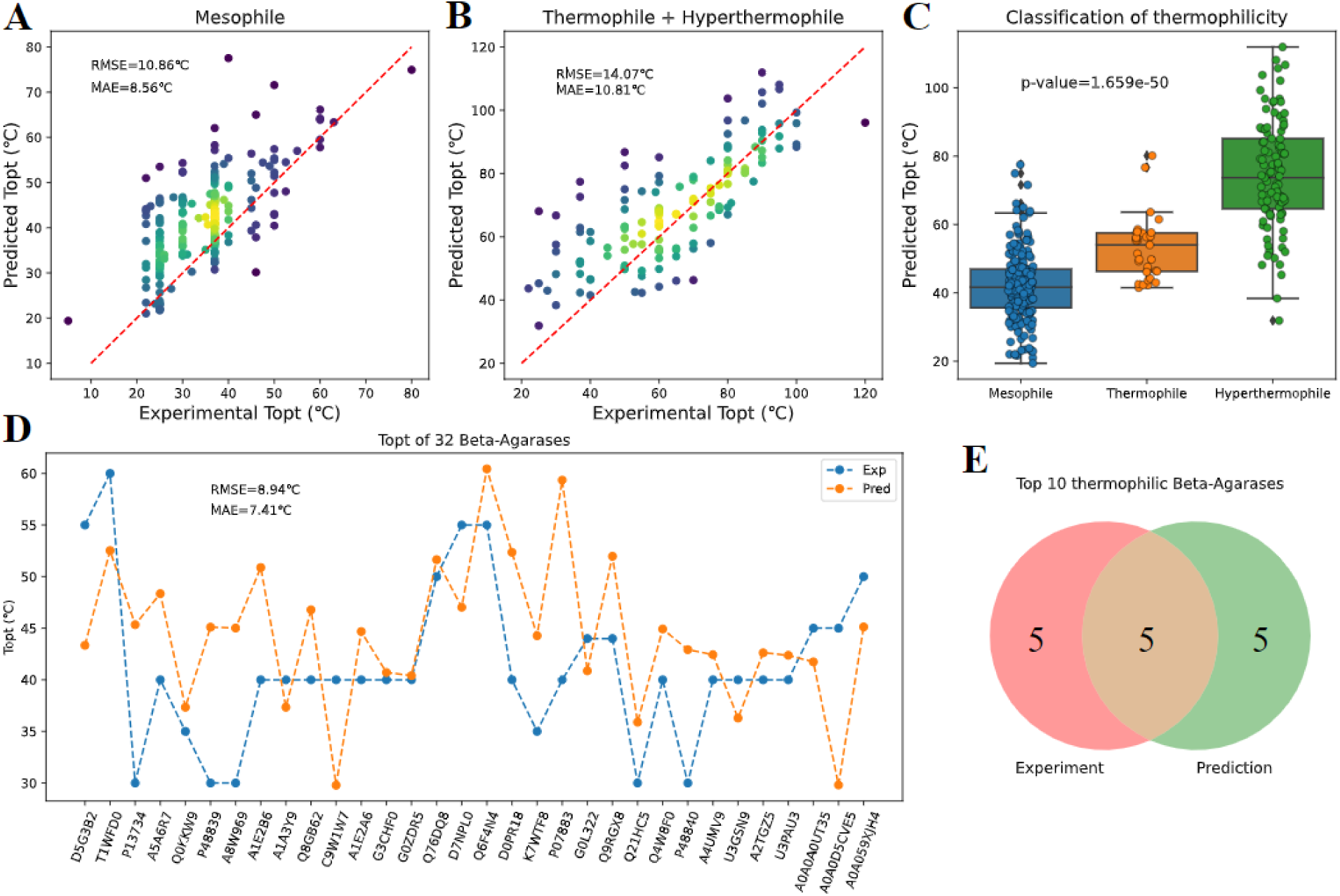
The performance of Seq2Topt on microorganisms and enzymes of different thermophilicities. (A) Experimental and predicted *T*_*opt*_ values of enzymes from mesophilic microorganisms (RMSE=10.86℃ and MAE=8.56℃). (B) Experimental and predicted *T*_*opt*_ values of enzymes from thermophilic and hyperthermophilic microorganisms (RMSE=14.07℃ and MAE=10.81℃). (C) The distributions of predicted *T*_*opt*_ values of enzymes from mesophilic, thermophilic, and hyperthermophilic microorganisms (p-value<0.001). Blue: mesophile; Orange: thermophile; Green: hyperthermophile. (D) Experimental and predicted *T*_*opt*_ values of 32 Beta-Agarases (RMSE=8.94℃ and MAE=7.41℃). Blue: experimental measurement; Orange: prediction. (E) The venn diagram of top 10 thermophilic Beta-Agarases determined by experiments and predicted by Seq2Topt.

Next, Seq2Topt was used to select thermophilic beta-agarases with the experimental data obtained from Chu et al., 2016 [7]. The overall prediction error of Seq2Topt on 32 beta-agarases was relatively low (**Figure 3D**), in comparison to the prediction error of Preoptem on the holdout test set (**section 3.1**). The 50% overlap between predicted and experimental top 10 thermophilic beta-agarases suggested that Seq2Topt, though trained on a dataset of general enzymes instead of a restricted scope, could identify thermophilic enzymes based on protein sequences (**Figure 3E**).

### 3.3 Deciphering the key sequence information for enzyme optimal temperature

To investigate how residue attention weights capture important sequence information for enzyme *T*_*opt*_, 4 methyltransferases (EC 2.1.1.) and 4 glycosidases (EC 3.2.1.) from mesophilic, thermophilic and hyperthermophilic microorganisms were selected from 328 enzymes in the previous section (**section 3.2**) to analyze the spatial distribution of residue attention weights and protein structural features (e.g., helix) (**see SI, Table S2**). For methyltransferases, the enrichment of local maximums of attention weights in helical regions can be observed (**Figure 4A-D**), especially for Ribosomal RNA small subunit methyltransferase G of *Escherichia coli* (**Figure 4A**) and rRNA adenine N-6-methyltransferase of Bacillus subtilis (**Figure 4D**).

**Figure 4.**
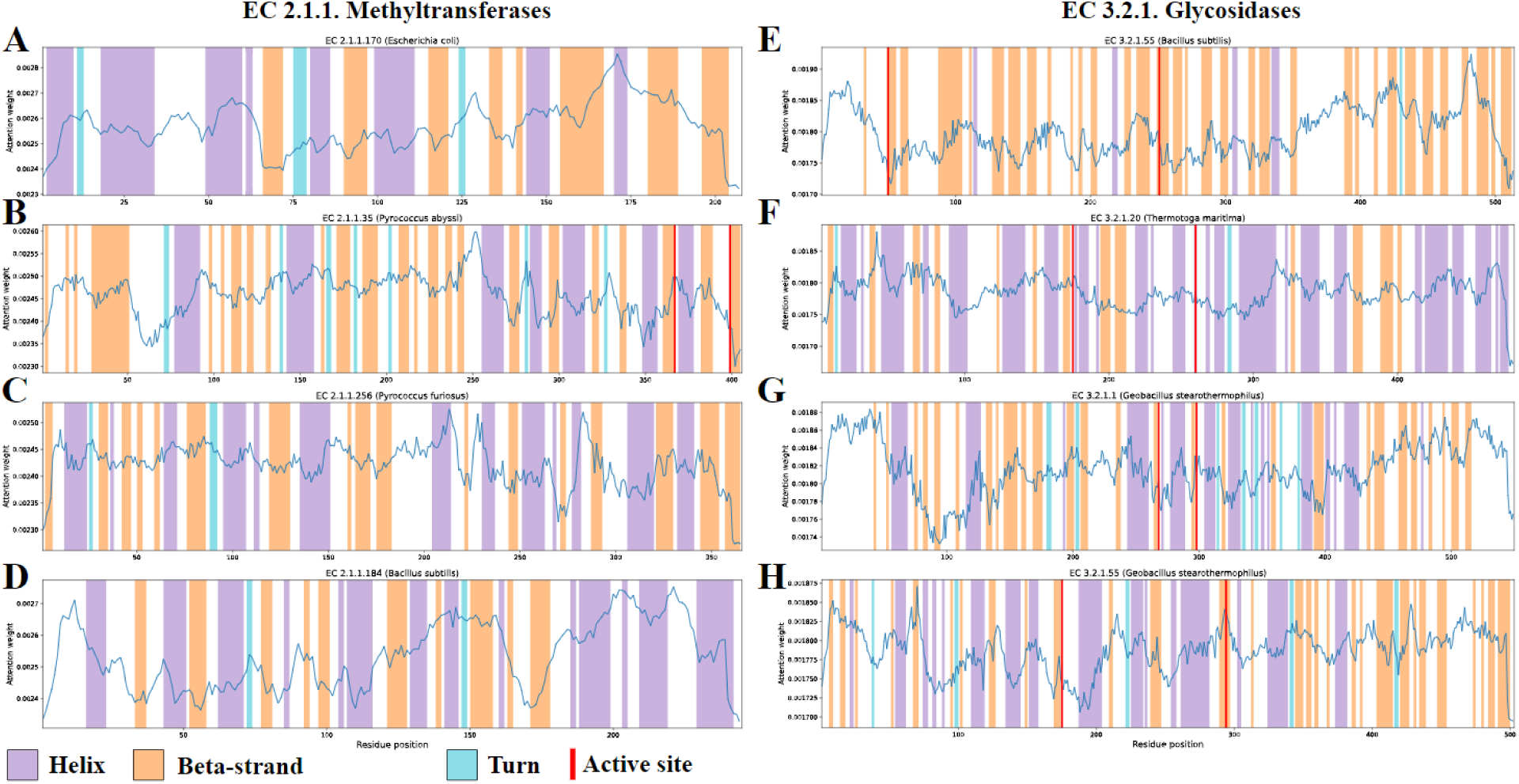
The spatial distributions of residue attention weights and protein structural features for (A) Ribosomal RNA small subunit methyltransferase G of *Escherichia coli* (P0A6U5, EC2.1.1.170), (B) tRNA (uracil(54)-C(5))-methyltransferase of *Pyrococcus abyssi* (Q9UZR7, EC 2.1.1.35), (C) tRNA (guanine(6)-N2)-methyltransferase of *Pyrococcus furiosus* (Q8U248, EC 2.1.1.256), (D) rRNA adenine N-6-methyltransferase of *Bacillus subtilis* (P13956, EC 2.1.1.184), (E) Arabinoxylan arabinofuranohydrolase of *Bacillus subtilis* (Q45071, EC 3.2.1.55), (F) Alpha-glucosidase of *Thermotoga maritima* (O33830, EC 3.2.1.20), (G) Alpha-amylase of *Geobacillus stearothermophilus* (P06279, EC 3.2.1.1), and (H) Intracellular exo-alpha-(1->5)-L-arabinofuranosidase of Geobacillus stearothermophilus (Q9XBQ3, EC 3.2.1.55). Purple: Helix; Orange: Beta-strand; Blue: Turn; Red line: Active site.

Regarding glycosidases, the analysis found that local maximums of attention weights concentrate in beta-strand regions (**Figure 4E-H**), especially for Arabinoxylan arabinofuranohydrolase of *Bacillus subtilis* (**Figure 4E**) and Alpha-amylase of *Geobacillus stearothermophilus* (**Figure 4G**). The helical regions of methyltransferases and beta-strand regions of glycosidases have both been found important for enzyme thermoactivity [17,18]. Furthermore, the local maximums of attention weights at residue 367 of tRNA (uracil(54)-C(5))-methyltransferase of *Pyrococcus abyssi* (**Figure 4B**) and residue 294 of Intracellular exo-alpha-(1->5)-L-arabinofuranosidase of *Geobacillus stearothermophilus* (**Figure 4H**) showed that Seq2Topt could capture the importance of active sites for enzyme *T*_*opt*_.

### 3.4 Prediction of the shift of enzyme optimal temperature caused by point mutations

This study used Seq2Topt to predict *T*_*opt*_ values of wild-types and mutants for xylose isomerases (XIs) of *Thermoanaerobacterium thermosulfurigenes* (TT) and *Thermotoga neapolitana* (TN) [19], beta-glucosidase (BGL) of *Trichoderma reesei* (TR) [20], and sucrose phosphorylase (SP) of *Bifidobacterium breve* (Bbr) [21] (**see SI, Table S3**). The experimental data of those enzymes was not included in the training set of Seq2Topt. The prediction accuracy of *T*_*opt*_ values of wild-type and mutated enzymes was examined, and RMSE, MAE scores were 8.24℃ and 6.37℃, respectively (**Figure 5A**). The xylose isomerases of *Thermoanaerobacterium thermosulfurigenes* (TT_XI) were not included in further analysis, due to low prediction accuracy **(see SI, Figure S4**). For the sucrose phosphorylase of *Bifidobacterium breve* (Bbr_SP), Seq2Topt qualitatively predicted that P134C/L343F and L341V/L343F could decrease and increase the *T*_*opt*_, respectively (**Figure 5B**). For the beta-glucosidase of *Trichoderma reesei* (TR_BGL), Seq2Topt identified that the mutation of L167W could enhance the *T*_*opt*_, but failed to predict the increase of *T*_*opt*_ caused by P172L/F250A (**Figure 5C**). For the xylose isomerase of *Thermotoga neapolitana* (TN_XI), Seq2Topt successfully predicted that P59Q and P63A mutations could decrease the *T*_*opt*_, and the combination of those two point mutations could result in a larger decrease of the *T*_*opt*_ than single point mutations (**Figure 5D**).

**Figure 5.**
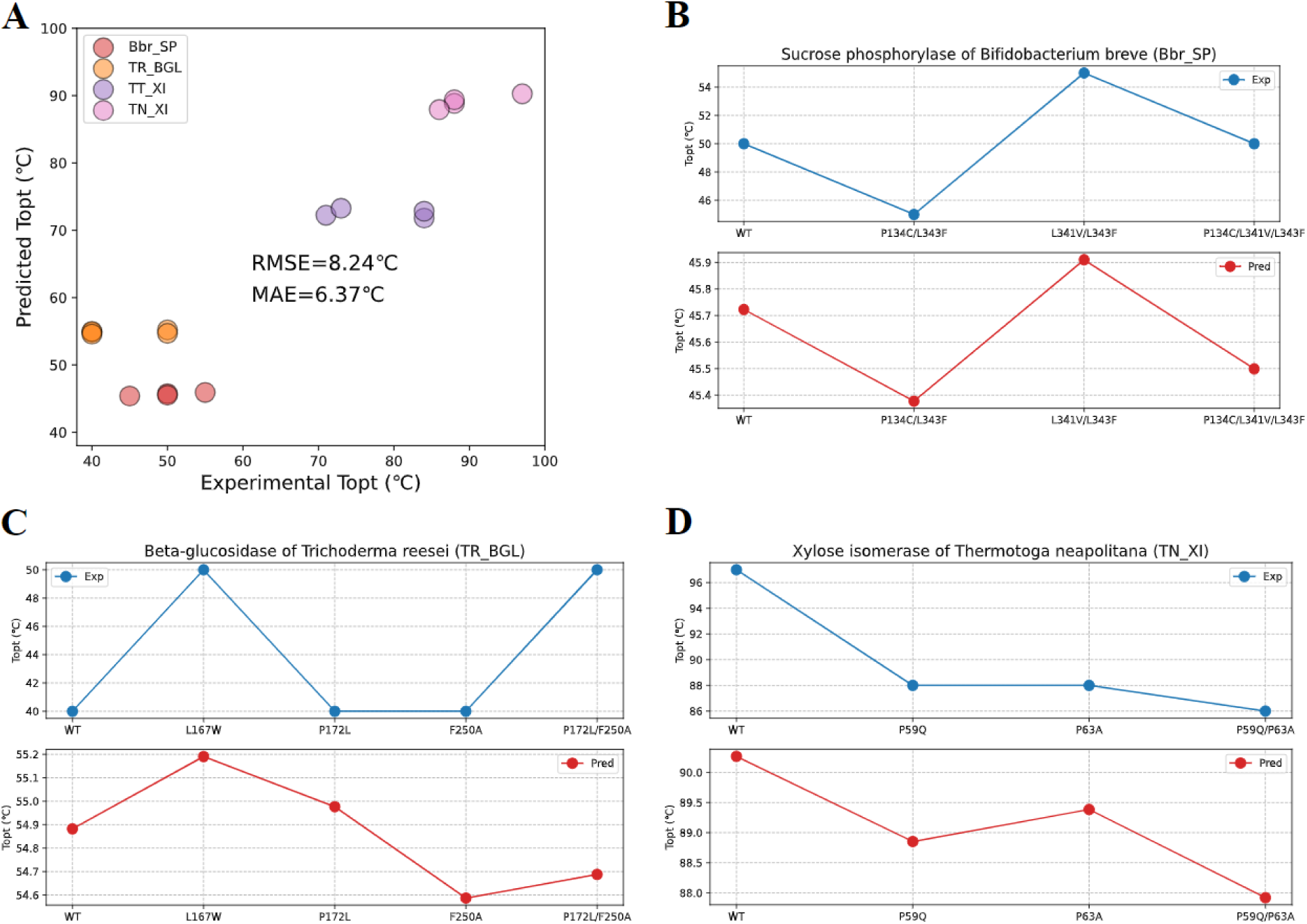
The prediction of enzyme optimal temperature shifts caused by mutations. (A) Experimental and predicted *T*_*opt*_ values of wild-types and mutants of 4 different enzymes (RMSE=8.24℃, MAE=6.37℃). (B) Experimental and predicted *T*_*opt*_ values of the wild-type and mutants of the sucrose phosphorylase of *Bifidobacterium breve* (Bbr_SP). (C) Experimental and predicted *T*_*opt*_ values of the wild-type and mutants of the beta-glucosidase of *Trichoderma reesei* (TR_BGL). (D) Experimental and predicted *T*_*opt*_ values of the wild-type and mutants of the xylose isomerase of *Thermotoga neapolitana* (TN_XI). TT: *Thermoanaerobacterium thermosulfurigenes*; TN:*Thermotoga neapolitana*; TR: *Trichoderma reesei*; HC: *Heyndrickxia coagulans*; BGL: beta-glucosidase; SP: sucrose phosphorylase; XI: xylose isomerase.

### 3.5 Seq2pHopt and Seq2Tm: use protein sequences to predict enzyme optimal pH and melting temperature

The model architecture of Seq2Topt (**Figure 1**) was used to construct predictive models of enzyme optimal pH (*pH*_*opt*_) and melting temperature (*T*_*m*_). This study developed Seq2pHopt for *pH*_*opt*_ and Seq2Tm for *T*_*m*_ with the default hyperparameters (**section 2.3**),and both of them used the protein sequence as the only input. In the training process, the RMSE scores of Seq2Tm and Seq2pHopt were reduced from around 16℃ to 7.57℃ and from 2 to 0.92, respectively (**Figure 6AD**). For MAE and R2 scores in the training process, please **see SI, Figure S5,6**. Seq2Tm and Seq2pHopt both achieved good accuracy on held-out test sets used by DeepTM [11] and EpHod [12], which are best models of *T*_*m*_ and *pH*_*opt*_ released (**Figure 6BE**). In comparison to DeepTM that requires both the protein sequence and OGT as inputs, Seq2Tm could reach a closely good prediction accuracy without using the OGT (**Figure 6C**).

**Figure 6.**
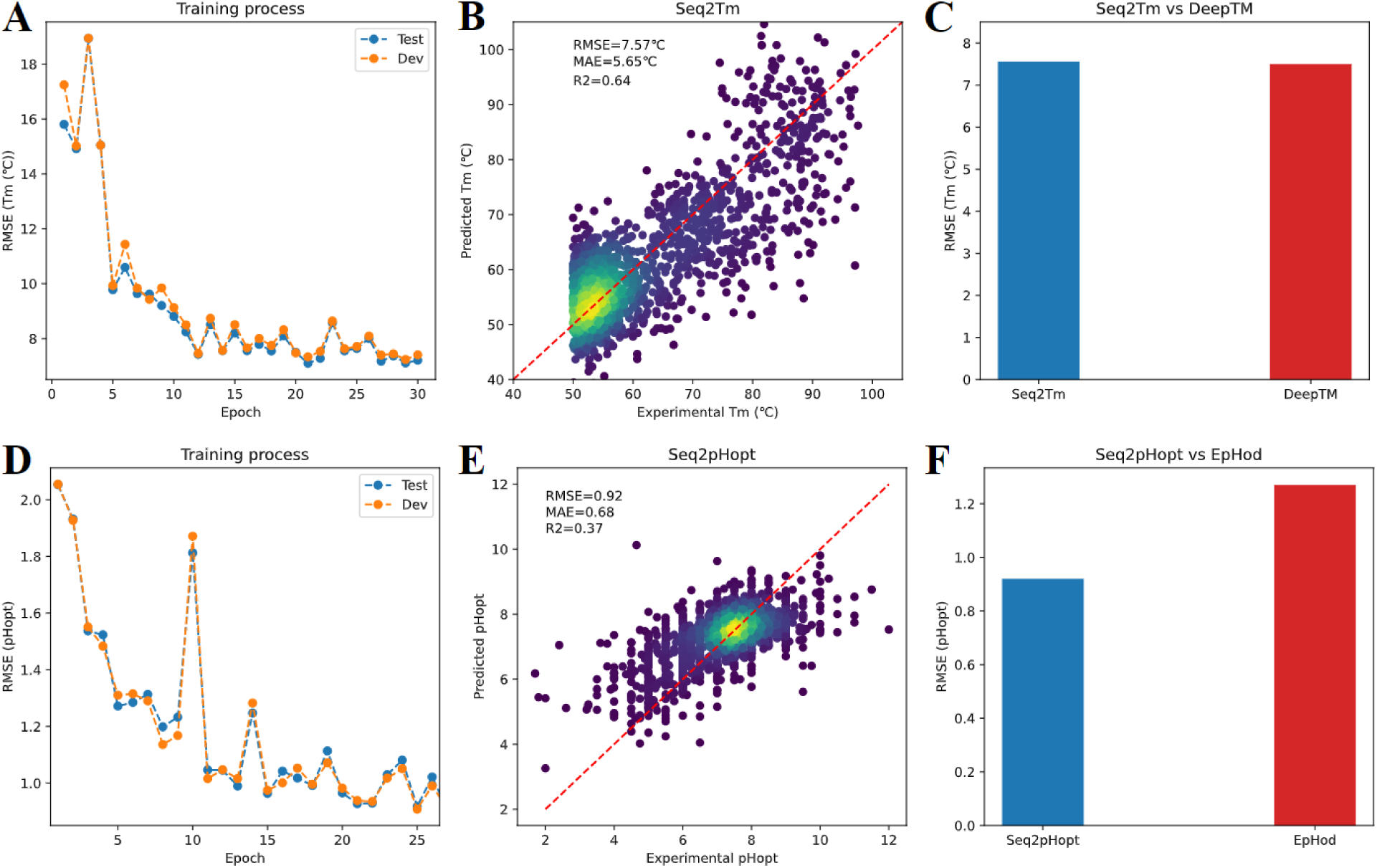
The assessment of prediction accuracy for Seq2Tm and Seq2pHopt. (A) The RMSE scores of *T*_*m*_ prediction during the training process. (B) Experimental and predicted *T*_*m*_ by Seq2Tm (RMSE=7.57℃, MAE=5.65℃, and R2=0.64). (C) The comparison of RMSE scores of *T*_*m*_ prediction by Seq2Tm and DeepTM (RMSE of Seq2Tm=7.57℃ and RMSE of DeepTM=7.5℃). (D) The RMSE scores of *pH*_*opt*_ prediction during the training process. (E) Experimental and predicted *pH*_*opt*_ by Seq2pHopt (RMSE=0.92, MAE=0.68, and R2=0.37). (F) The comparison of RMSE scores of *pH*_*opt*_ prediction by Seq2pHopt and EpHod (RMSE of Seq2pHopt =0.92 and RMSE of EpHod=1.27).

For *pH*_*opt*_ prediction, Seq2pHopt outperformed EpHod with a lower RMSE score (**Figure 6F**). The RMSE scores of DeepTM and EpHod were reported in their articles [11][12]. Generally speaking, Seq2Tm and Seq2pHopt had superior prediction accuracy on *T*_*m*_ and *pH*_*opt*_, respectively.

## 4. Discussion

The gap of experimental data and expensive cost of enzyme *T*_*opt*_ demand an accurate and easy-to-use predictive model, and this study managed to tackle this challenging task and developed Seq2Topt, a deep learning model that can predict enzyme *T*_*opt*_ just from protein sequences. Three main elements of Seq2Topt were ESM-2 embedding generation, multi-head attention, and residual dense neural networks (**Figure 1**). Compared with conventional feature extraction methods of protein sequences, such as one-hot encoding [10] or k-mer based dictionary embedding [22,23], ESM-2, as a pre-trained large language model of proteins, has the advantage of being able to learn the information of structures and functions hidden in sequences [13]. In contrast to single-head attention, multi-head attention could improve the prediction accuracy by focusing on different parts of protein sequences simultaneously [24] and effectually capturing important regions (**section 3.3**), which has been demonstrated by the higher accuracy of Seq2pHopt than EpHod that uses single-head attention [12] (**Figure 6F**). In addition, the use of residue dense neural networks instead of multiple linear layers could effectively reduce the vanishing and exploding gradient issues in deep neural networks [25]. Also, oversampling on entries at the high temperature value range, to some extent, compensated for the imbalanced distribution of enzyme *T*_*opt*_ values in the dataset (**section 2.1**). As a result, Seq2Topt outperformed the other existing enzyme *T*_*opt*_ predictor just using protein sequences, Preoptem [10], with RMSE close to 10℃ and R2 close to 0.5.

Case studies of selecting thermophilic beta-agarases (**section 3.2**) and predicting enzyme *T*_*opt*_ shifts caused by point mutations (**section 3.4**) manifested that Seq2Topt could be applied to enzyme mining and computational design of enzymes via fast screening the effect of mutations. Expectedly, the combination of Seq2Topt and generative deep learning might lead to predictor-guided generator optimization [26] of enzymes, enabling automatic enzyme design. Furthermore, all three accurate predictive models of enzyme *T*_*opt*_, *pH*_*opt*_, and *T*_*m*_ can potentially improve the performance of condition dependent enzyme *k*_*cat*_ prediction (e.g., DLTKcat [23] or MPEK [27]) by informing the catalytic optimum.

Despite the achievement of Seq2Topt, there still exist some limitations for Seq2Topt, which hinder the improvement of prediction accuracy. First, its accuracy in the high temperature value range is relatively low (**Figure 2**), and the main reason lies in the imbalance of the dataset. Also, the size of the dataset used to develop Seq2Topt is much smaller than those of other deep learning models for proteins, such as the dataset of EpHod containing 9855 enzymes [12]. One possible solution is to append the dataset of enzyme *T*_*opt*_ by curating more entries from databases (e.g., BRENDA [2]) or conducting high-throughput enzyme assays, especially for the high temperature value range. Another shortcoming of Seq2Topt is that it cannot account for the impact of environmental factors on the thermoactivity of enzymes [28], such as pH [29] or enzyme concentrations in assays [30]. Including metadata of curated experimental measurements might resolve such shortcoming.

In conclusion, Seq2Topt is an accurate and easy to use (the only input needed is the protein sequence) deep learning predictor of enzyme *T*_*opt*_, in spite of some limitations discussed above. As envisaged, Seq2Topt can potentially accelerate enzyme discovery for desired properties from “biological dark matter” and enzyme engineering with *in-silico* design, and might give rise to a powerful prediction platform of enzymes.

## Key points

● Seq2Topt can accurately predict enzyme optimal temperature values just from protein sequences.
● Seq2Topt can predict the shift of enzyme optimal temperature caused by point mutations.
● Residue attention weights of Seq2Topt can reveal important sequence regions for enzyme thermoactivity.
● The architecture of Seq2Topt can be used to build predictors of other enzyme properties (e.g., optimal pH).

## Acknowledgements

The authors would like to acknowledge the use of the University of Oxford Advanced Research Computing (ARC) facility (http://dx.doi.org/10.5281/zenodo.22558) in carrying out this work. The authors would like to thank Simiao Zhao and Yishun Lu for offering technical advice.

## Author contributions

Sizhe Qiu constructed the deep learning model and performed case studies. Bozhen Hu assisted in model construction and contributed to model optimization. Sizhe Qiu and Bozhen Hu together produced the first draft. Jing Zhao and Weiren Xu assisted in case studies and reviewed the manuscript. Aidong Yang supervised this research project and critically reviewed the manuscript.

## Data availability statement

The code and data are openly available at https://github.com/SizheQiu/Seq2Topt.

## Abbreviation

Bbr: Bifidobacterium breve
BGL: beta-glucosidase
CNN: convolutional neural network Leaky
ReLU: leaky rectified linear unit
MAE: mean absolute error
MSE: mean squared error
MUT: mutant
OGT: optimal growth temperature
*pH*_*opt*_: optimal pH
R2: r-squared, the coefficient of determination
RMSE: root mean squared error
SP: sucrose phosphorylase
*T*_*m*_: melting temperature
*T*_*opt*_: optimal temperature
TN: Thermotoga neapolitana (TN)
TR: Trichoderma reesei
TT: Thermoanaerobacterium thermosulfurigenes
WT: wild type
XI: xylose isomerase

