## Supplementary material for "Seq2Topt: a sequence-based deep learning predictor of enzyme optimal temperature": Figure S1-6, Table S1-3

### Authorship

Sizhe Qiu^1^, Bozhen Hu^2,3^, Jing Zhao^4,5^, Weiren Xu^5^, Aidong Yang^1^*

^1^Department of Engineering Science, University of Oxford, OX1 3PJ, United Kingdom

^2^AI Division, School of Engineering, Westlake University, Hangzhou 310030, China

^3^Zhejiang University, Hangzhou 310058, China

^4^State Key Laboratory of Biocatalysis and Enzyme Engineering, Hubei Collaborative Innovation Center for Green Transformation of Bio-Resources, Hubei Key Laboratory of Industrial Biotechnology, School of Life Sciences, Hubei University, Wuhan 430062, China

^5^Tianjin Institute of Pharmaceutical Research Co., Ltd., Tianjin 300301, China

### 1. Supplementary methods

#### 1.1 Software and code availability

All scripts were written in python. The deep learning model was implemented using PyTorch v1.7.1. The computer used in this work was a Dell Latitude Laptop with intel core i7 CPU. The model was trained with GPU RTX8000 provided by Advanced Research Computing (ARC) service in the University of Oxford [[1]](https://paperpile.com/c/rih2ej/McUo). Figures were edited using InkScape (<https://inkscape.org/>). The code and data used to generate results of this paper are available at <https://github.com/SizheQiu/Seq2Topt>.

#### 1.2 Evaluation metrics

To quantitatively assess the prediction accuracy, R2 (Eq. S1), RMSE (Eq. S2), and MAE (Eq. S3) were computed for each test.

$$R^{2}=\frac{\sum_{i=1}^{n} (y_{ie}-y_{ip})^{2}}{\sum_{i=1}^{n} (y_{ie}-\underline{y})^{2}} (Eq. S1)$$

$$RMSE=\sqrt{\frac{1}{n}\sum_{i=1}^{n} (y_{ie}-y_{ip})^{2}} (Eq. S2)$$

$MAE = \frac{1}{n}\sum_{i=1}^{n} \left| y_{ie}-y_{ip} \right| (Eq. S3)$

### 2. Supplementary figures

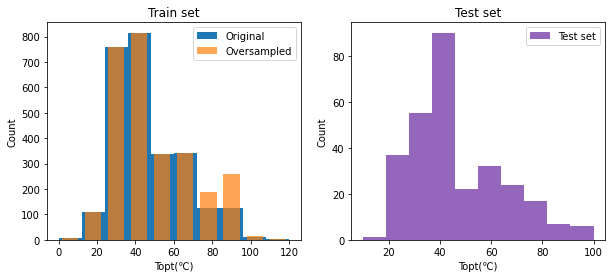

Figure S1. The distribution of $T_{opt}$ values in training and test sets.

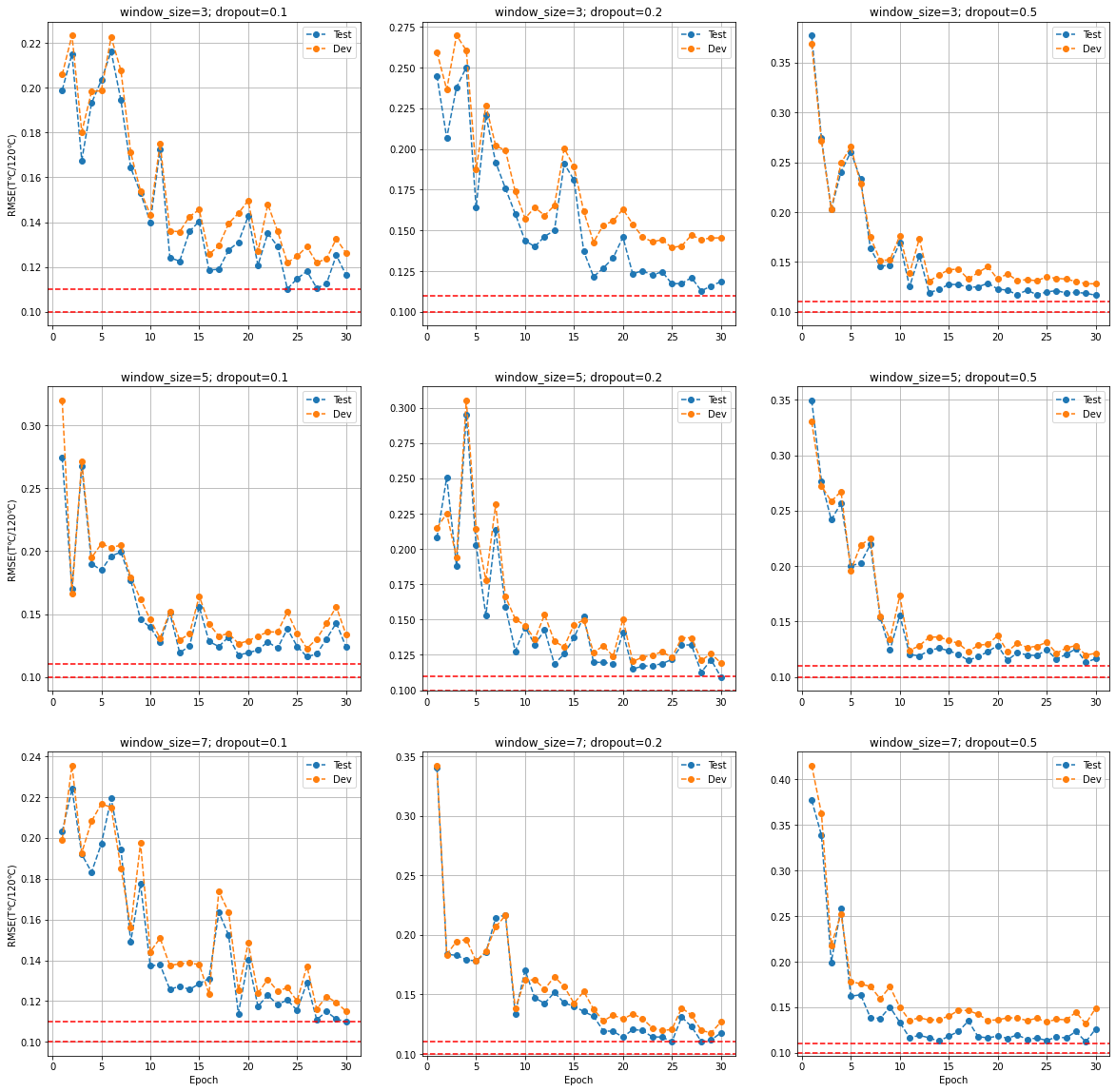

Figure S2. Hyperparameter optimization for $T_{opt}$ prediction on the sliding window size of the CNN and the dropout rate. The optimal set of hyperparameters is {window size:5, dropout rate: 0.2, number of attention heads: 4, number of residual dense blocks: 3}.

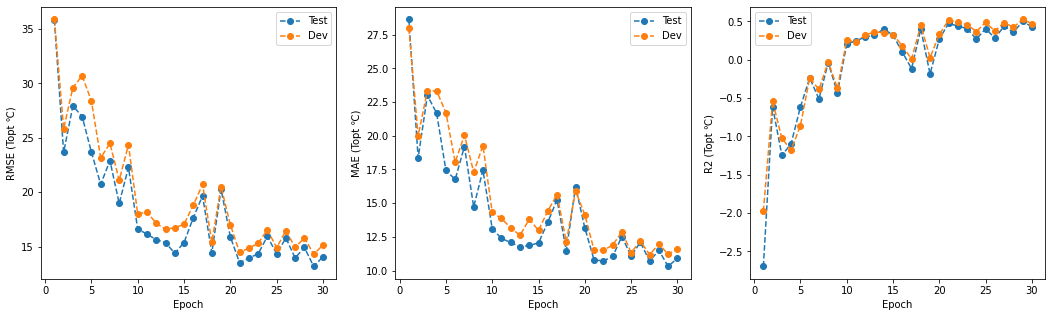

Figure S3. The training process of Seq2Topt in 30 epochs. Topt: optimal temperature.

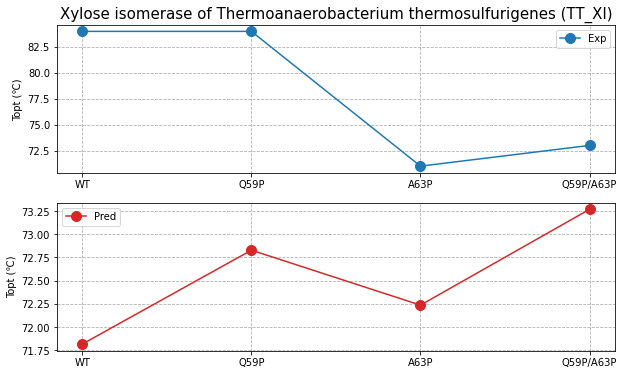

Figure S4. Experimental and predicted $T_{opt}$ values of the wild-type and mutants of the xylose isomerase of *Thermoanaerobacterium thermosulfurigenes* (TN_XI).

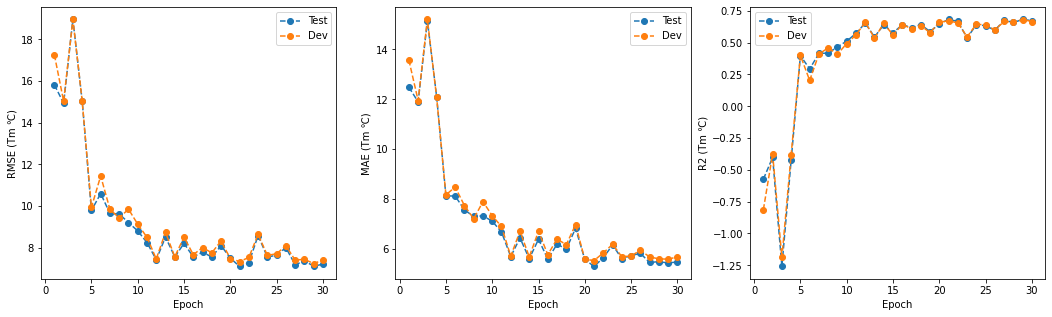

Figure S5. The training process of Seq2Tm in 30 epochs. Tm: melting temperature.

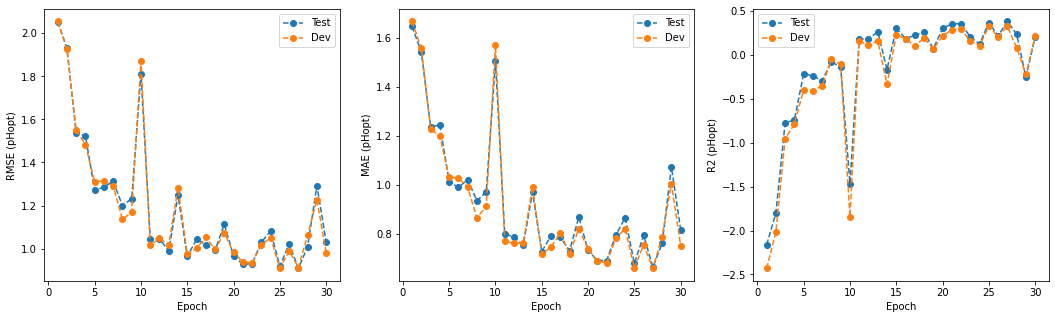

Figure S6. The training process of Seq2pHopt in 30 epochs. pHopt: optimal pH.

### 3. Supplementary tables

Table S1. Selected microorganisms for the estimation of thermophilicity

| Organism | Class | OGT (℃) |
| --- | --- | --- |
| Escherichia coli | Mesophile | 36 |
| Bacillus subtilis | Mesophile | 30 |
| Staphylococcus aureus | Mesophile | 37 |
| Geobacillus stearothermophilus | Thermophile | 54 |
| Geobacillus thermoleovorans | Thermophile | 60 |
| Geobacillus thermodenitrificans | Thermophile | 57 |
| Pyrococcus furiosus | Hyperthermophile | 96 |
| Thermotoga maritima | Hyperthermophile | 75 |
| Pyrococcus abyssi | Hyperthermophile | 90 |
| Aquifex aeolicus | Hyperthermophile | 80 |

Table S2. Selected enzymes for the analysis of residue attention weights.

| Uniprot ID | Organism | Class | EC | Topt (℃) |
| --- | --- | --- | --- | --- |
| P0A6U5 | Escherichia coli | Mesophile | 2.1.1.170 | 37 |
| Q9UZR7 | Pyrococcus abyssi | Hyperthermophile | 2.1.1.35 | 50 |
| Q8U248 | Pyrococcus furiosus | Hyperthermophile | 2.1.1.256 | 70 |
| P13956 | Bacillus subtilis | Mesophile | 2.1.1.184 | 31 |
| Q45071 | Bacillus subtilis | Mesophile | 3.2.1.55 | 45 |
| O33830 | Thermotoga maritima | Hyperthermophile | 3.2.1.20 | 90 |
| P06279 | Geobacillus stearothermophilus | Thermophile | 3.2.1.1 | 80 |
| Q9XBQ3 | Geobacillus stearothermophilus | Thermophile | 3.2.1.55 | 40 |

Table S3. Information of wild-type and mutated enzymes used in case studies.

| Enzyme | Organism | Mutation | Topt (℃) | Reference |
| --- | --- | --- | --- | --- |
| Beta-glucosidase | Trichoderma reesei | WT | 40 | [[2]](https://paperpile.com/c/rih2ej/0uKAT) |
|  |  | L167W | 50 |  |
|  |  | P172L | 40 |  |
|  |  | F250A | 40 |  |
|  |  | P172L/F250A | 50 |  |
| Xylose isomerase | Thermotoga neapolitana | WT | 97 | [[3]](https://paperpile.com/c/rih2ej/aeJSD) |
|  |  | P59Q | 88 |  |
|  |  | P63A | 88 |  |
|  |  | P59Q/P63A | 86 |  |
| Xylose isomerase | Thermoanaerobacterium thermosulfurigenes | WT | 84 | [[3]](https://paperpile.com/c/rih2ej/aeJSD) |
|  |  | Q59P | 84 |  |
|  |  | A63P | 71 |  |
|  |  | Q59P/A63P | 73 |  |
| Sucrose phosphorylase | Bifidobacterium breve | WT | 50 | [[4]](https://paperpile.com/c/rih2ej/C87r) |
|  |  | P134C/L343F | 45 |  |
|  |  | L341V/L343F | 55 |  |
|  |  | P134C/L341V/L343F | 50 |  |

* values were extracted from plots using WebPlotDigitizer (<https://github.com/automeris-io/WebPlotDigitizer>) [[5]](https://paperpile.com/c/rih2ej/5QeY).
